# Anthropomorphic left ventricular mesh phantom: a framework to investigate the accuracy of SQUEEZ using Coherent Point Drift for the detection of regional wall motion abnormalities

**DOI:** 10.1101/840595

**Authors:** Ashish Manohar, Gabrielle Colvert, Andrew Schluchter, Francisco Contijoch, Elliot R. McVeigh

## Abstract

We present an anthropomorphically accurate left ventricular (LV) phantom derived from human CT data to serve as the ground truth for the optimization and the spatial resolution quantification of a CT-derived regional strain metric (SQUEEZ) for the detection of regional wall motion abnormalities. Displacements were applied to the mesh points of a clinically derived end-diastolic LV mesh to create analytical end-systolic poses with physiologically accurate endocardial strains. Normal function as well as regional dysfunction of four sizes (1, 2/3, 1/2, and 1/3 AHA (American Heart Association) segments as core diameter), each exhibiting hypokinesia (70% reduction in strain) and subtle hypokinesia (40% reduction in strain), were simulated. Regional shortening (RS_CT_) estimates were obtained by registering the end-diastolic mesh to each simulated end-systolic mesh condition using a non-rigid registration algorithm. Ground-truth models of normal function and of hypokinesia were used to identify the optimal parameters in the registration algorithm, and to measure the accuracy of detecting regional dysfunction of varying sizes and severities. For normal LV function, RS_CT_ values in all 16 AHA segments were accurate to within ±5%. For cases with regional dysfunction, the errors in RS_CT_ around the dysfunctional region increased with decreasing size of dysfunctional tissue.

## 1. Introduction

The assessment of regional cardiac function has implications in the diagnosis, treatment, and follow up of patients with cardiac diseases such as myocardial ischemia^1, 2^, heart failure^3^, cardiotoxicity^4^, and dyssynchrony^5, 6^. Left-ventricular (LV) ejection fraction (EF), global longitudinal strain (GLS), and other global metrics have been useful in the assessment of overall LV function, but fail to provide information on the regional health of the myocardium. Additionally, significant reduction in global function metrics appear only in advanced disease stages whereas regional dysfunction often precedes global dysfunction.

The current gold-standard for the non-invasive assessment of regional LV function is cardiovascular magnetic resonance (CMR) tagging^7–10^. However, CMR requires extended breath holds, acquisition over multiple heart beats, and manual contouring^7, 11^. The growing number of patients with metallic medical device implants further limits the clinical use of CMR.

Recent advances in x-ray computed tomography (CT) have made possible the acquisition of an entire 3D volume of the heart from a single table position in ∼140 ms, which implies a series of functional images spanning the full cardiac cycle can be obtained within a single heartbeat^12–16^. Additionally, the spatial resolution (0.4 x 0.4 x 0.6 mm^3^ nominal voxel size) allows for the detection and tracking of the fine endocardial texture comprising trabeculae carneae and papillary muscles^17^. The drawback of CT is patient exposure to ionizing radiation; however, due to recent advancements in CT technology, especially in the last 5 years, the average dose received by a patient from a functional cardiac CT scan is ∼3 mSv, which is the average dose received from natural sources in a year^18^.

SQUEEZ^17^ is a new method introduced to measure regional endocardial strain from 4DCT images acquired with routine clinical protocols. SQUEEZ exploits the high fidelity of x-ray CT to track features of the endocardium, which are used by a non-rigid point set registration technique^19^ to derive displacements of points on the endocardium across the cardiac cycle. This displacement estimate is used to obtain information on the regional strain of the endocardium. SQUEEZ has shown to be capable of differentiating normal from dysfunctional myocardial regions in pigs^17^, has been compared with CMR tagging derived circumferential shortening in canines^20^, and more recently, baseline estimates of regional shortening (RS_CT_) values derived from SQUEEZ in the normal human LV were established^21^. In addition to the LV, SQUEEZ has shown to be capable of identifying regional differences in systolic function in complex right-ventricle (RV) geometries^22^.

Cross-modality comparisons of regional strain measures in humans is a difficult task and the measurements made maybe subject to modality-specific biases as each imaging modality has inherent limitations ranging from operator variability to through-plane motion. Additionally, trabeculation of the endocardium is species dependent, with human hearts having more trabeculation than either porcine or canine hearts; hence, the prior optimization of SQUEEZ will likely need adjustment for human hearts. Significant effort has been laid on the development of computerized phantoms of the human heart which provide an essential ground-truth in the optimization and evaluation of various imaging devices and techniques. Some of these phantoms are derived directly from patient image data (voxel phantoms)^23–26^, while others are defined using mathematical primitives^27, 28^. The voxel phantoms are anatomically accurate but are limited in their flexibility for modelling different motions and anatomies, while on the other hand, the mathematical phantoms offer a great deal of modelling flexibility but do not accurately represent the human anatomy. A new breed of hybrid phantoms that combine the advantages of the above two classes have been developed^29–32^. These phantoms have been very useful in optimizing and evaluating various imaging systems for optimal patient diagnosis and treatment.

In this paper, we describe the development of an anthropomorphically accurate LV mesh phantom with defined correspondences between endocardial points as a function of time across the cardiac cycle^33^. The phantom is derived from a high-resolution CT scan of a human LV and displacements were applied to each point to create analytical end systolic poses exhibiting physiologically accurate endocardial strains. Using the phantom, we tune the parameters of the registration algorithm to achieve optimal registration between the end diastolic and the end systolic LV poses. Additionally, we also quantify the spatial resolution limits of SQUEEZ for the estimation of regional wall motion abnormalities, thereby performing the first analytical evaluation of SQUEEZ using an anthropomorphically accurate phantom of the human LV.

## 2. Methods

### 2.1 Image Acquisition

The end diastolic phase of a human cardiac CT scan was used as the baseline image for phantom development. The image was acquired under IRB approved protocols at the National Institutes of Health (Bethesda, Maryland) using retrospective electrocardiogram (ECG) gating with inspiratory breath-hold. The scanner used was a 320 detector-row (Aquilion ONE, Canon Medical Systems, Otawara, Japan), which enabled complete coverage of the heart from a single table position. The scan parameters were: tube current - 280 mA, kVP - 100 kV, and short scan acquisition with a gantry revolution time of 275 ms. Full R-R image data spanning a complete cardiac cycle was acquired and the end diastolic phase was reconstructed at 0% of the R-R phase. Optimal opacification from contrast dye was achieved using real-time bolus tracking.

The image was reconstructed into a matrix of 512 x 512 x 320 voxels using the manufacturer’s (Canon Medical Systems) standard reconstruction algorithm (reconstruction diameter: 173.5 mm; kernel: Cardiac FC08), as implemented clinically. The image had voxel dimensions of 0.34 x 0.34 x 0.50 mm^3^ in the *x*, *y*, and *z* dimensions respectively. Neither the imaging parameters nor subject enrolment were determined by the authors for the purpose of our particular study. The images used in this work were directly obtained from the clinic with acquisition and reconstruction parameters determined by the clinical protocol used for that study.

### 2.2 Image Processing & Mesh Extraction

The LV endocardium was segmented with ITK-SNAP^34^ using the active contour region growing module (thresholding type = high pass; threshold = 450; smoothness = 10; seed radius = 5; all other parameters were set as default). All further processing steps were performed in MATLAB (MathWorks Inc.).

The binary LV segmentation was loaded into MATLAB in its native orientation (original axial slices) and resolution. Prior to extracting the endocardial mesh, the binary LV volume was resampled to an isotropic resolution of 0.5 x 0.5 x 0.5 mm^3^ by linear interpolation. The isotropic binary volume was rotated sequentially about the *x* and *y* axes to align the LV long axis with the *z* axis; hence, the longitudinal direction of the model is equivalent to the *z*-direction.

The binary LV volume was subsampled to a resolution of 2 x 2 x 2 mm^3^ by retaining every fourth (2mm/0.5mm) voxel in *x*, *y*, and *z* directions to achieve a spatial sampling consistent with previously published SQUEEZ analyses^17, 20–22^. A mesh of the boundary between the LV blood pool and the endocardium was extracted using the *isosurface* built-in MATLAB routine. The mesh comprised of 5257 faces and 2662 mesh points. A summary of the process is shown in Fig. 1A-C.

**Fig. 1.**
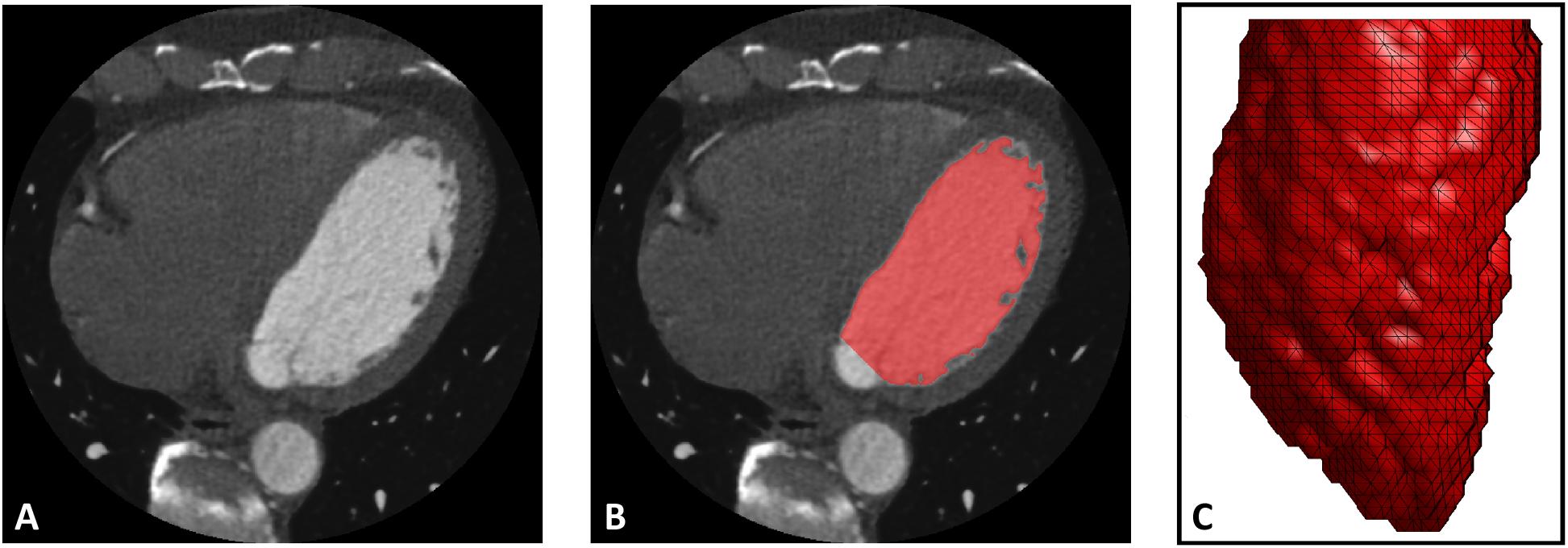
Image segmentation and mesh extraction. (**A**) Axial slice of the CT image. (**B**) Axial slice with the LV blood volume segmentation overlaid in red (with the mitral valve and the LV outflow-tract defined as the cut-off points). (**C**) 3-D rendering of the extracted LV endocardial mesh, looking at the lateral wall with the anterior wall to the left, and the inferior wall to the right.

### 2.3 Strain Model

The end diastolic mesh, obtained from the LV segmentation as described in Sec 2.2, was deformed to an analytical end systolic pose by prescribing displacements to the mesh points to achieve physiologically accurate end systolic strains. The three deformations modeled were:

1. Longitudinal strain (ε_zz_): constant −21%^35^ from apex to base
2. Circumferential strain (ε_cc_): linear reduction from −44% at the apex to −34% at the base^35^
3. Azimuthal rotation around the long axis (Δθ): linear decrease^36^ in rotation from 13° in the counter-clockwise direction at the apex to 6.9° in the clockwise direction at the base, when viewed from apex to base^37^

Azimuthal rotation represents the rotation of the LV endocardium during systole; changes in Δθ as a function of long axis position create torsion in the LV model. The EF of the model was 70%, which was consistent with CT-based LV EFs measured in normal hearts^21^. Figure 2 shows the peak end systolic strain values as a function of *z*-position along the long axis of the LV.

**Fig 2.**
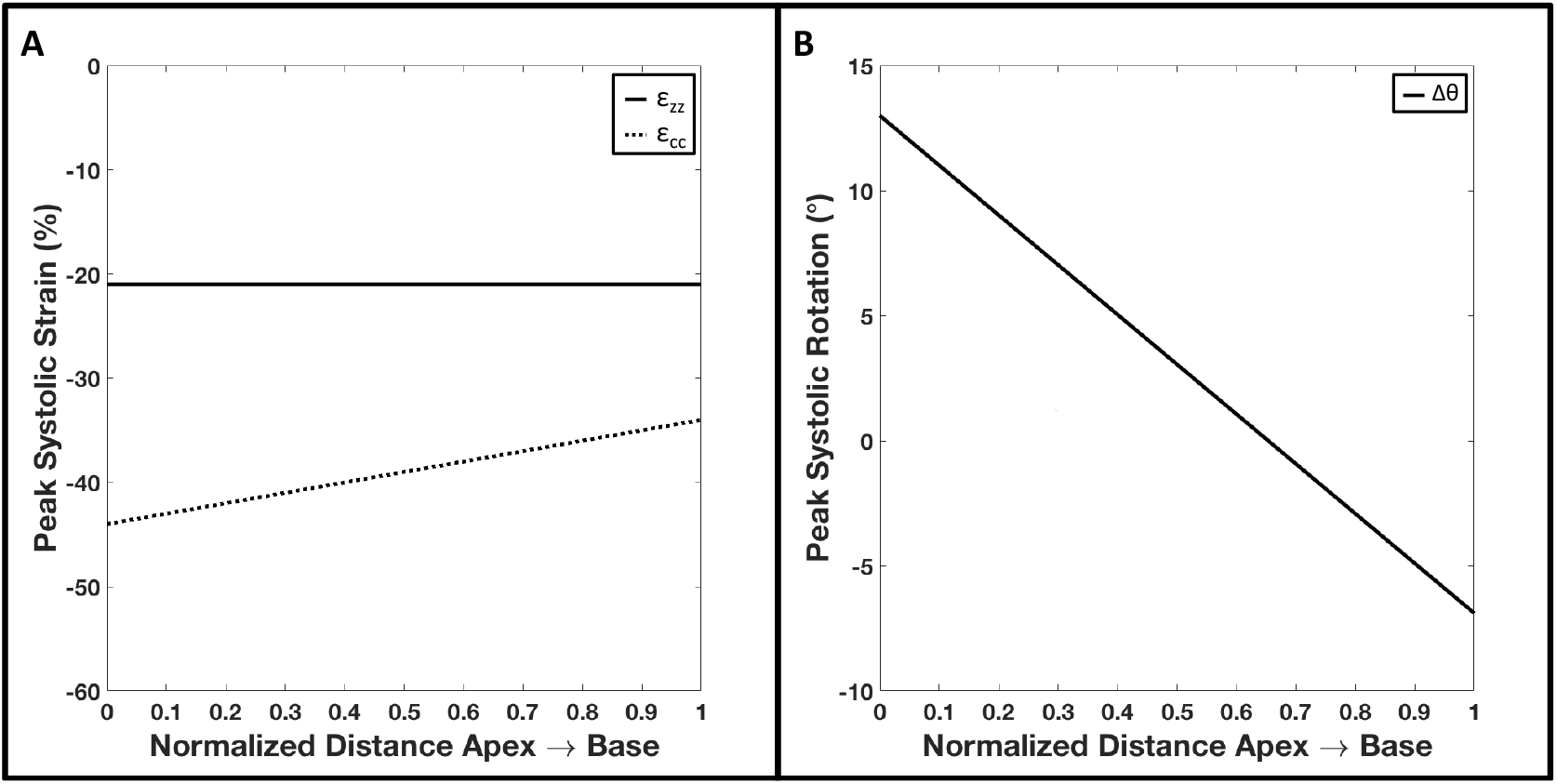
Components of the peak end systolic strain model as a function of *z*-position along the long axis of the LV from apex (position 0) to base (position 1). (**A**) Peak end systolic longitudinal (ε_zz_; solid line) and circumferential (ε_cc_; dotted line) strains as a function of the LV long axis (**B**) Peak end systolic LV rotation as a function of the LV long axis (positive rotation is counter-clockwise when viewed from apex looking up towards the base).

#### 2.3.1 Displacement functions

Displacements were prescribed to each mesh point (*x,y,z*) in order to attain the desired three deformations specified above according to the following equations

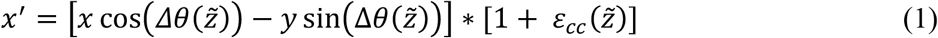

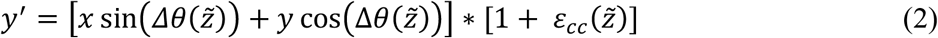

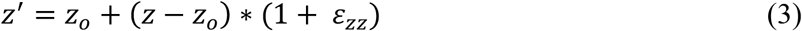

where (*x’,y’,z’*) are the new displaced coordinates, 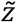 is the normalized *z* coordinate (0 at the apex and 1 at the base), and *z_o_* is the fixed apical *z* coordinate.

#### 2.3.2 Regional dysfunction model

Regional dysfunction was incorporated into the strain model by prescribing a reduction factor in the ε_zz_ and the ε_cc_ components over a particular region of interest. The dysfunction model had three adjustable parameters:

1. Position: spatial coordinates of the center of the dysfunctional region
2. Severity: reduction factor in strain at the center of the dysfunctional region
3. Size: diameter at which a 13% reduction from the desired dysfunctional severity occurred; defined as the ‘core’ diameter

To ensure continuity of the LV tissue, a 2D sigmoid function was used to define the dysfunctional region to create a smooth transition radially outward from the center (where severity is maximum) to the normal tissue (see Appendix A). The sizes of the dysfunctional regions were reported in terms of fractions of an AHA segment^38^ (corresponding to the American Heart Association’s 17 segment model) defining the core of the dysfunctional region. The extent of the core was defined as the diameter at which a 13% reduction from the desired dysfunctional severity occurred; this corresponded to the flat ‘table-top’ region of the sigmoid function. A representative human end diastolic LV circumference of 170 mm was used as reference to define the AHA segments. For example, 1/2 an AHA segment translated to a core of 14 mm (170 mm/12).

Two severities of dysfunction were investigated in this paper: 1) hypokinesia, defined by a peak 70% reduction at the center in the ε_zz_ and the ε_cc_ components and 2) subtle hypokinesia, defined by a peak 40% reduction at the center in the ε_zz_ and the ε_cc_ components.

#### 2.3.3 Texture smoothing

To mimic the CT image sampling of the end systolic pose of the ventricle more accurately, a low-pass filter was applied on the mesh using the *iso2mesh*^39^ MATLAB toolbox (method = ‘lowpass’; iterations = 4; alpha = 0.4). This was motivated by the change in LV endocardial texture during systole^40, 41^ due to collapsing spaces between trabeculae coupled with spatial resolution limitations of the CT scanner; as the trabeculae come closer together in the analytical phantom due to myocardial contraction, the spatial frequency of the endocardial surface increases. Due to the scanner’s finite resolution (point spread function), these high frequency structures can no longer be resolved giving rise to a smoothed texture of the endocardium at end systole. To simulate this texture change of the LV, a filter was applied. The filter was a Laplace filter applied in two steps; a forward step (smoothing; eqn. 4) and a backward step (volume conserving; eqn. 5) according to the following equations^42^

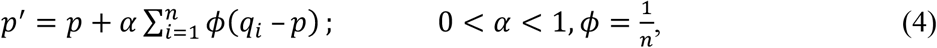

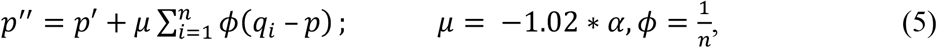

where *p*, *p’*, and *p’’* are the original, intermediate, and final positions of a mesh point *p* and *i = 1,2,3,…,n* is the index number of the *n* neighboring mesh points *q.* The parameters *α*, *μ*, and number of iterations were chosen such that the frequency response of the applied filter was similar to that of the Standard reconstruction kernel of a 256 detector-row clinical scanner (Revolution CT, General Electric Medical Systems, Wisconsin; see Appendix B). Figure 3 shows the end diastolic (Fig. 3A) and the analytically derived final end systolic poses (with smoothing) under normal function (Fig. 3B) and a sample case exhibiting regional dysfunction (Fig. 3C).

**Fig. 3.**
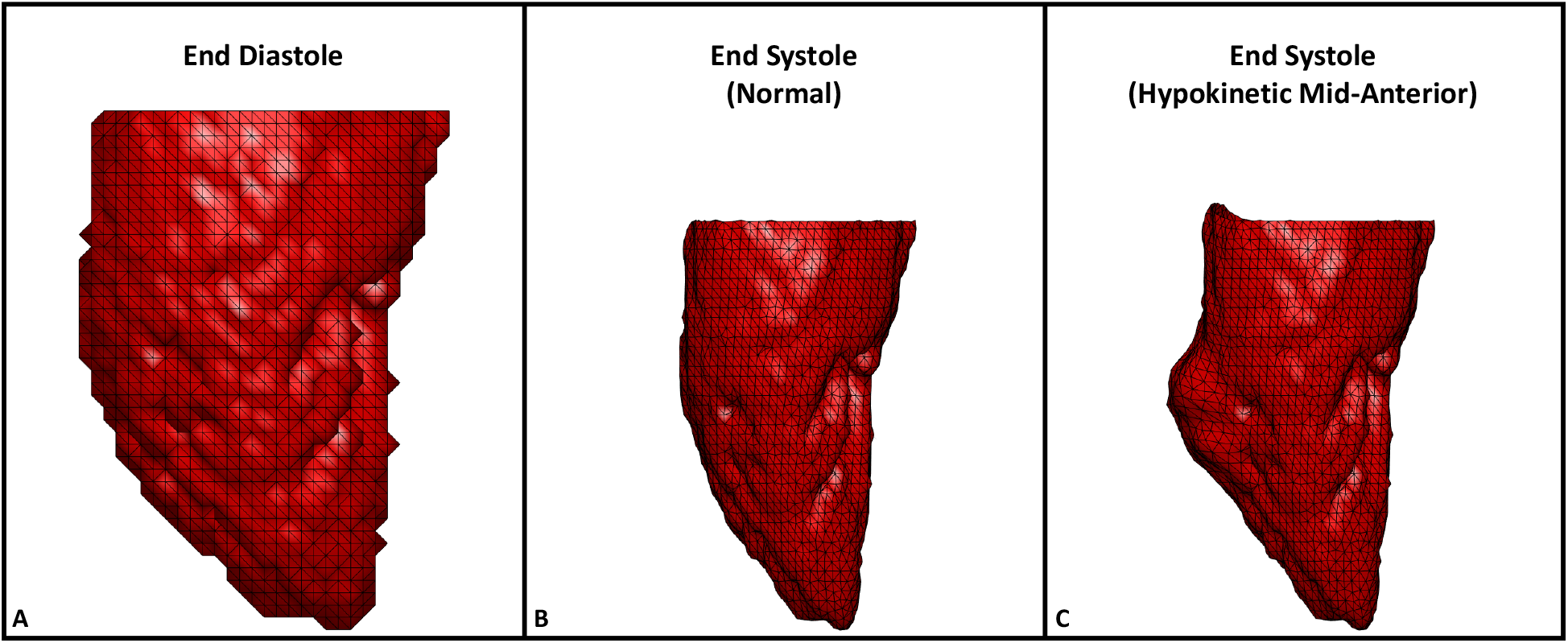
End diastolic and analytically derived end systolic LV poses. (**A**) End diastolic mesh derived from human LV clinical CT data. (**B**) Analytically derived end systolic pose with normal function. (**C**) Analytically derived end systolic pose exhibiting regional hypokinesia of 1/2 an AHA segment (14 mm) as core diameter at the mid-anterior segment of the LV with normal strain values in all other segments. All three views are of the LV lateral wall with the anterior wall to the left and the inferior wall to the right.

### 2.4 Non-rigid Point Set Registration

Coherent Point Drift (CPD), a probabilistic non-rigid point set registration technique^19^, was used to register the end diastolic template mesh to the end systolic target mesh. Since our model had a perfect 1:1 correspondence, the registration accuracy could be validated with the known ground-truth. However, in the clinical scenario, a 1:1 correspondence does not exist between the end diastolic and the end systolic mesh points. From 10 4DCT scans of human LVs with normal function (CT measured LV EF: 73% ± 4%), meshes were extracted at end diastole and end systole and the ratio of the number of mesh points at end systole to end diastole was calculated to be 46% ± 7%; therefore, only 50% of the mesh points (every alternate point) in the analytically derived end systolic poses were retained.

Additionally, a uniform distribution of random noise in the range of ± 0.6 mm was added to the coordinates (*x*,*y*,*z*) of the down-sampled end systolic pose to make the noise independent. The limits of the added noise were determined by calculating the natural variability in the detection of the endocardial edge for different contrast-to-noise levels obtained by changing the x-ray tube current settings (see Appendix C). This down-sampled end systolic pose with added noise was used as the target for the registration of the template end diastolic mesh.

#### 2.4.1 Identification of optimal set of CPD parameters

The CPD algorithm has 2 free parameters: *β*, and *λ*. *β* is the width of the Gaussian smoothing kernel and *λ* is the regularization weight^19^. Both *β* and *λ* define the coherence of motion between neighboring points; an increase in either value, forces the points to move in a more coherent manner, while lower values allow for more localized deformations.

For our application of detecting a subtle wall motion abnormality, we needed the registration algorithm to balance global coherent motion with the ability to track localized deformations. The warping of the template end diastolic mesh to the target end systolic mesh had to be performed in a physiologically accurate manner (coherent motion), while simultaneously capturing regional abnormalities in cardiac contraction (localized changes in the expected motions). From visual inspection trials, performed on phantoms representing both normal LV function and regional hypokinetic function (1/2 AHA segment (14 mm) as core diameter), we observed *β* values between [0.5, 2] in steps of 0.1 and *λ* values between [1, 25] in steps of 1 covered the domain of values of interest. The above identified values for *β* and *λ* defined 400 (16*β* x 25*λ*) sets of possible parameters for the registration algorithm. From these 400, we sought to identify one set of parameters which yielded the ‘best’ (defined in Sec. 2.4.2) registration between end diastole and end systole under both, normal and regional dysfunction conditions. The 1/2 AHA segment hypokinetic model was chosen based on two factors: 1) from visual inspection of wall motion abnormalities seen on CT images, this was considered smaller than the average size of dysfunction and 2) by optimizing for hypokinetic function, the registration fit would be better capable of capturing subtle wall motion abnormalities. The core of the 1/2 AHA segment hypokinetic region contained 36 mesh points.

#### 2.4.2 Statistical evaluation

The accuracy and quality of the registration fit was assessed based on four metrics:

1. Global *l*^2^ norm: The global *l*^2^ norm was computed for all mesh points of the LV according to the formula

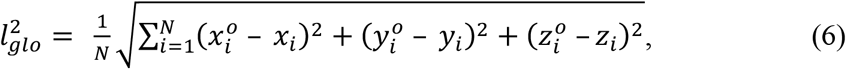

where *(x°,y°,z°)* are the coordinates of the ground-truth mesh, *(x,y,z)* are the corresponding coordinates of the registered mesh, and *i = 1,2,3,…,2662(N)* is the index of the mesh points.
2. Regional *l*^2^ norm: The regional *l*^2^ norm was computed over the mesh points of the programmed dysfunctional region of the LV according to the formula

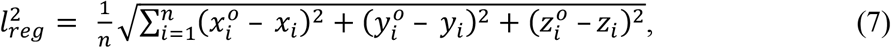

where *(x°,y°,z°)* are the coordinates of the ground-truth mesh, *(x,y,z)* are the corresponding coordinates of the registered mesh, and *i = 1,2,3,…,320(n)* is the index of the mesh points within the programmed dysfunctional region.
3. Euclidean distance: Unlike the *l*^2^ norm which provides an average measure of goodness of fit for all points in consideration, Euclidean distance was calculated on a point by point basis according to the formula

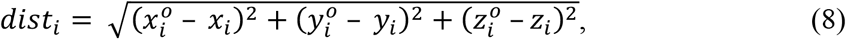

where *(x°,y°,z°)* are the coordinates of the ground-truth mesh, *(x,y,z)* are the corresponding coordinates of the registered mesh, and *i = 1,2,3,…,2662(N)* is the index of the mesh points.
4. SQUEEZ: SQUEEZ^17^ is a user-independent method developed to measure endocardial regional shortening (RS_CT_; it is the geometric mean of circumferential and longitudinal strain). It is measured by tracking features on the endocardium from 4DCT data and is computed as the square root of the ratio of areas of corresponding patches between a template mesh and a target mesh, according to the formula

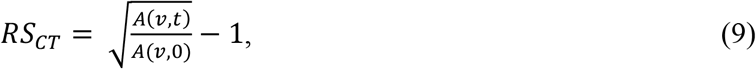

where *A(v,t)* refers to the area of a patch *v* at time *t* and *A(v,0)* is the area of the same patch at time *t = 0* (end diastole). Consistent with previously published SQUEEZ analyses, each patch was defined as a single triangular face of the end diastolic template mesh; the average patch size at end diastole was 2.6 ± 0.5 mm^2^. Recently, from a cohort of 25 subjects with normal LV function, McVeigh et al. determined the average peak end systolic RS_CT_ value over the entire LV endocardium to be −34% ± 5%^21^.

Only those values of *β* and *λ*, from the pool of 400, that yielded registration fits that satisfied the following inclusion criteria were shortlisted:

1. Global *l*^2^ norm in the normal case ≤ 0.015 (bottom 90% of all *l*^2^ norms; errors were consistently low for the simulated normal LV function)
2. Regional *l*^2^ norm in the dysfunctional case ≤ 0.07 (bottom 40% of all regional *l*^2^ norms)
3. 95% of RS_CT_ values in the normal case to be within ± 5%^21^ from the ground-truth
4. 90% of mesh points (2396 in number) in both the normal and the dysfunctional case to be within ± 1 mesh point distance (± 2 mm) from the ground-truth
5. RS_CT_ values in all 16 AHA segments in the normal case to be within ± 5% from the ground-truth
6. RS_CT_ values in 15 normal AHA segments (excluding the mid-anterior segment containing the programmed dysfunctional region; AHA segment 7) in the dysfunctional case to be within ± 5% from the ground-truth
7. Average RS_CT_ value in abnormal AHA segment 7 (mid-anterior segment containing the programmed dysfunctional region) in the dysfunctional case ≥ −26% (average ground-truth RS_CT_ value in the normal LV = −31%, hence cut-off threshold for abnormal function is - 31% + 5% = −26%). The theoretical RS_CT_ value in the core of the hypokinetic region was −12 ± 2.2%.

### 2.5 Quantifying Spatial Resolution Limits of SQUEEZ for the Detection of Regional Dysfunction

Four sizes of regional dysfunction (1, 2/3, 1/2, and 1/3 AHA segments translating to core diameters of 28, 19, 14, and 9.5 mm respectively), each modeled with hypokinesia and subtle hypokinesia (defined in Sec. 2.3.4) were used to sample the threshold of detection of abnormal strain with SQUEEZ. The optimal set of CPD parameters, as determined from Sec. 2.4.2, was used and the target end systolic meshes for each size and severity of dysfunction were prepared as outlined in Sec. 2.4 (down-sampled and independent noise added).

Box plots of the distribution of RS_CT_ values within the core of the dysfunctional regions, for both the ground-truth and the registered meshes, were used to quantify the accuracy. The spatial resolution limit of SQUEEZ under both regional hypokinetic and subtle hypokinetic functions was defined as the smallest lesion size that SQUEEZ can accurately detect to within ± 5% of the theoretical RS_CT_ value in the core of the dysfunctional regions.

## 3. Results

The results are presented under two sections: 1) identification of the optimal set of parameters for CPD registration and 2) quantification of the spatial resolution limits for the detection of regional wall motion abnormalities.

### 3.1 Optimal Set of Parameters for CPD Registration

From the 400 sets of parameters, only 6 sets of parameters satisfied the 7 inclusion criteria outlined in Sec. 2.4.2. *β* values only between 1 and 1.1 satisfied the inclusion criteria; high values of *β* imposed strong coherence of motion and hence, regional displacements (hypokinetic/infarct regions) were compromised while very low values of *β* led to hyper-flexible meshes. The values of *λ* ranged from 7 to 9 for *β = 1.1* and from 10 to 12 for *β = 1*. Also observed was an inter-dependency between *β* and *λ*; as the value of *β* (width of the Gaussian smoothing kernel) decreased, in order to maintain the overall quality of the mesh (coherence of point displacements), the value of *λ* (the regularization weight) had to increase.

Figure 4 plots the *l*^2^ norms as a function of *β* and *λ* under normal and regional dysfunction cases within the domain of *β* and *λ* values that satisfied the inclusion criteria. As seen from Fig. 4, the errors in both regional and global fits were stable and similar across all sets of shortlisted parameters (highlighted by the red boundaries). While any of the 6 shortlisted sets of parameters could be used, we needed to choose one operating point to proceed with the investigation of the spatial resolution of SQUEEZ for the detection of regional wall motion abnormalities. From Fig. 4B, for a value of *β = 1.1*, the errors across a wide range of *λ* values between [7,13] were stable and consistently low; this served as motivation to choose *β = 1.1* over *β = 1.0*. Among the three available *λ* values of 7, 8, and 9 for *β = 1.1*, *λ = 9* was chosen as the operating point to provide the highest regularization from among the shortlisted parameters. While our simulation was performed under controlled circumstances, human LVs artifactually lose features during systole in a non-coherent fashion (such as trabeculae collapsing and papillary muscles merging with the endocardium); hence for clinical LV scans, we expect that a higher regularization of the registration fit would decrease the influence of the artifactual non-rigid deformation caused by the above listed artifacts.

**Fig. 4.**
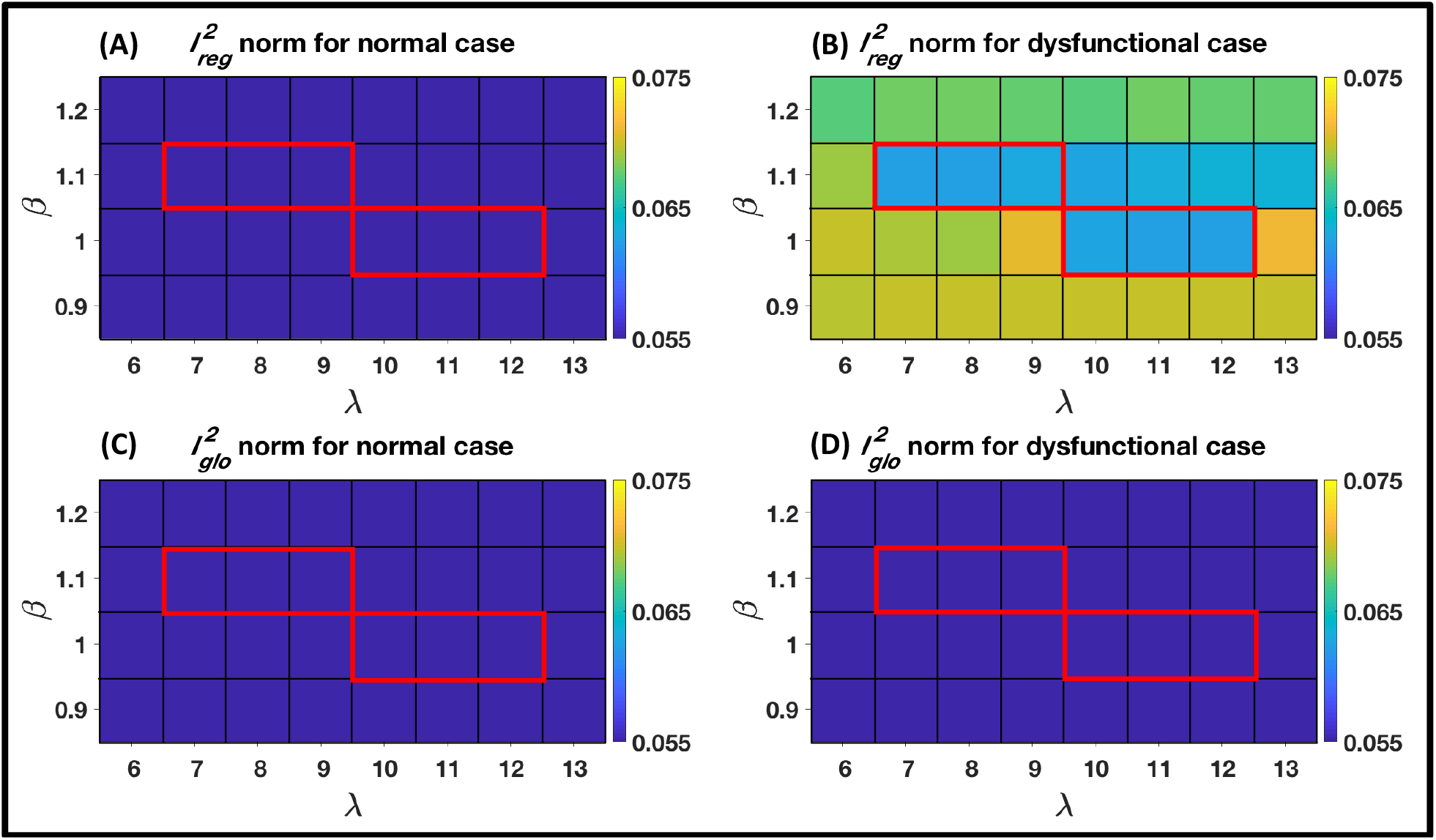
Registration errors as a function of *β* and *λ* within the domain of *β* and *λ* values that satisfied the inclusion criteria. (**A**) *l*^2^ norm computed over a region of the endocardium in the model with normal LV function that corresponded to the hypokinetic region in the regional dysfunction model. (**B**) *l*^2^ norm computed over the hypokinetic region in the regional dysfunction model. (**C**) *l*^2^ norm computed over the entire endocardium in the model with normal LV function. (**D**) *l*^2^ norm computed over the entire endocardium in the regional dysfunction model. The registration errors were stable and similar across all parameter sets that satisfied the inclusion criteria (highlighted by the red boundaries).

### 3.2 Spatial Resolution limits of SQUEEZ for the Detection of Regional Dysfunction

Four sizes of regional dysfunction were simulated with each size exhibiting hypokinesia and subtle hypokinesia (see Sec. 2.5). Figure 5 summarizes the results of SQUEEZ for the detection of hypokinesia for the four sizes of simulated regional dysfunction. Panels A and B show the end systolic poses of the anterior wall of the LV with RS_CT_ values mapped onto the endocardial surface (panel A: programmed ground-truth RS_CT_ maps; panel B: RS_CT_ maps derived from SQUEEZ). Panel C shows boxplots (bottom and top edges of each box correspond to the 25^th^ and the 75^th^ percentiles and the central line indicates the median) of the ground-truth (in blue) and the SQUEEZ derived (in pink) RS_CT_ values within the core of each of the four dysfunctional regions. For a detection threshold of ± 5% from the theoretical RS_CT_ value within the core of the dysfunctional regions, Fig. 5 demonstrates the smallest lesion size that can be confidently measured with SQUEEZ processing. Hypokinetic regions of 1 and 2/3 AHA segments as core diameter were accurately measured while the severity of dysfunction in the 1/2 and 1/3 AHA segments was underestimated.

**Fig. 5.**
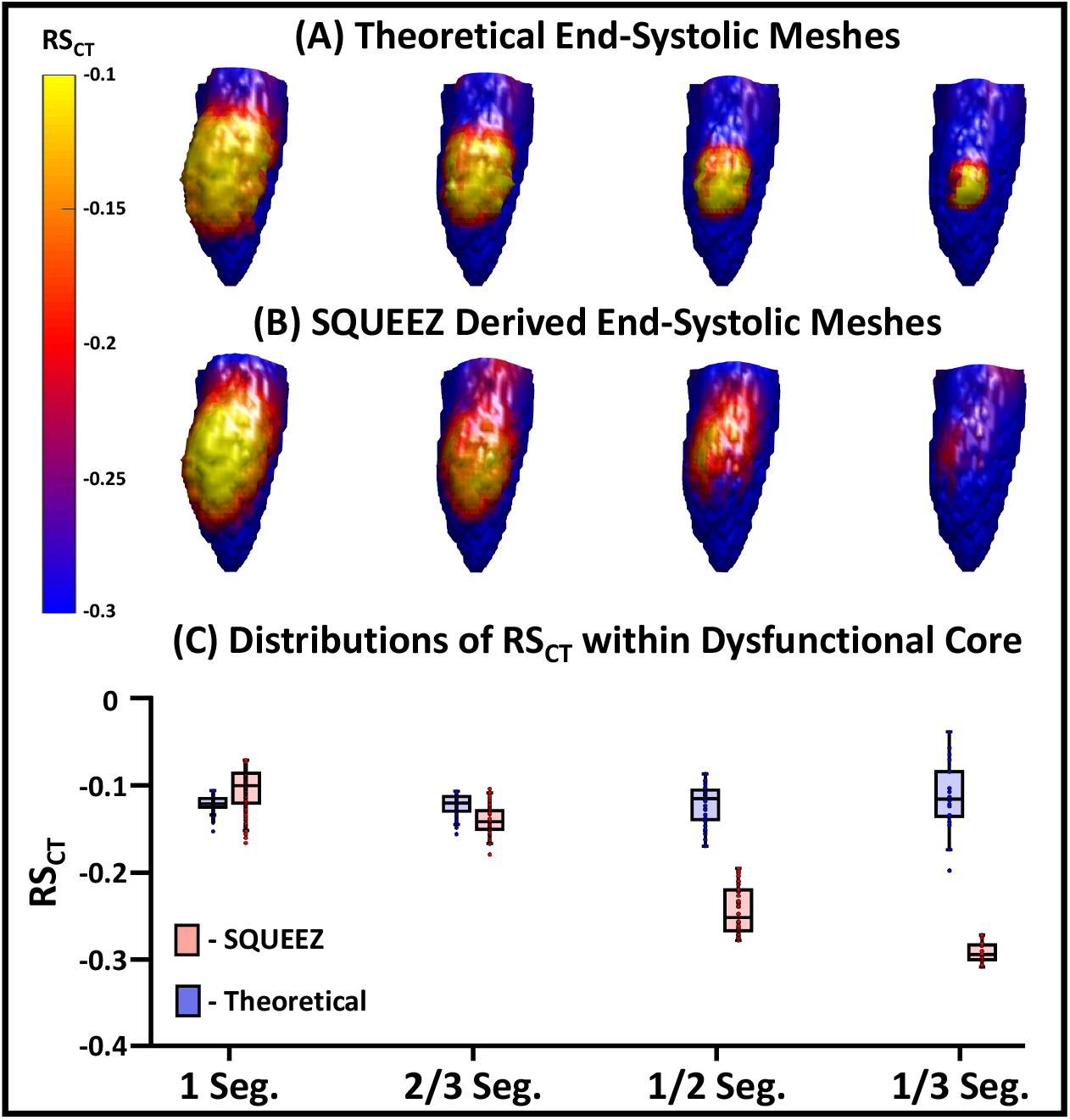
Detection of hypokinesia with SQUEEZ for four different sizes of regional dysfunction. (**A-B**) End systolic LV poses showing the anterior wall with regional shortening (RS_CT_) values mapped onto the endocardial surface for the four sizes of regional dysfunction (**A** – programmed ground-truth RS_CT_ maps; **B** – RS_CT_ maps obtained with SQUEEZ). Yellow corresponds to low function and blue corresponds to normal function. **(C)** Boxplots of ground-truth and SQUEEZ derived RS_CT_ values within the core of each of the four dysfunctional regions.

Similarly, Fig. 6 summarizes the results of SQUEEZ for the detection of subtle hypokinesia for the four sizes of simulated regional dysfunction. Panels A and B show the end systolic poses of the anterior wall of the LV with RS_CT_ values mapped onto the endocardial surface (panel A: programmed ground-truth RS_CT_ maps; panel B: RS_CT_ maps derived from SQUEEZ). Panel C shows boxplots of the ground-truth and the SQUEEZ derived RS_CT_ values within the core of each of the four dysfunctional regions. Again, subtle hypokinetic regions of 1 and 2/3 AHA segments as core diameters were accurately measured while the severity of dysfunction in the 1/2 and 1/3 AHA segments was underestimated.

**Fig. 6.**
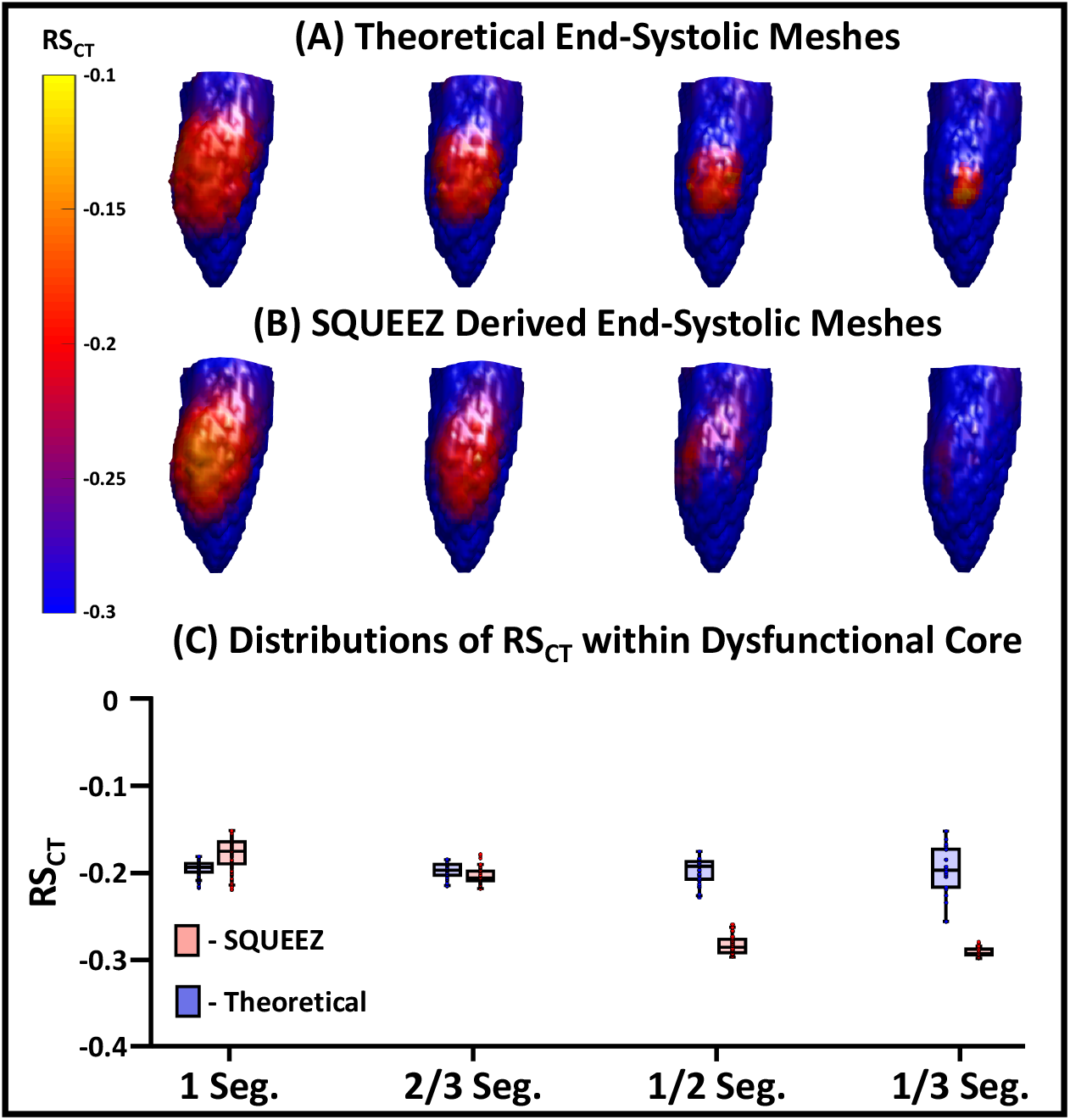
Detection of subtle hypokinesia with SQUEEZ for four different sizes of regional dysfunction. (**A-B**) End systolic LV poses showing the anterior wall with regional shortening (RS_CT_) values mapped onto the endocardial surface for the four sizes of regional dysfunction (**A** – programmed ground-truth RS_CT_ maps; **B** – RS_CT_ maps obtained with SQUEEZ). Yellow corresponds to low function and blue corresponds to normal function. **(C)** Boxplots of ground-truth and SQUEEZ derived RS_CT_ values within the core of each of the four dysfunctional regions.

As seen from Fig. 5 and 6, the reduction in strain in localized and subtle wall motion abnormalities was underestimated. While SQUEEZ was able to capture lesions of size greater than 2/3 AHA segments with high accuracy, this result suggested that secondary techniques need to be identified and developed to facilitate the accurate capture of smaller localized regions of dysfunction.

## 4. Discussion

This manuscript presents an anthropomorphic LV mesh phantom derived from human cardiac CT data and prescribed with physiologically accurate strain functions^35–37^ to create analytically known systolic poses. We have introduced an LV mesh model with programmable dysfunctional regions of various sizes, shapes, and severities for the first analytical evaluation of the spatial resolution of SQUEEZ for the human LV. A potential alternative to the work described here would be to compare strain estimates from SQUEEZ and an another imaging technique such as MR tagging. However, comparing strain values in the human heart measured with two different imaging techniques is fraught with problems; the human subject needs to be imaged with each technique in quick succession, and each technique has its own inherent limitations. The LV phantom developed in this work provides a platform to compare the measured strain values with the actual programmed ground-truth values. In addition to the measurement of regional shortening, the model also provides a platform to investigate the assessment of other LV phenomena such as dyssynchrony and endocardial twist, which were not examined in the work reported here. Another advantage of the developed mathematical phantom is that the analytically derived systolic poses can be used to create 3D printed physical phantoms, serving as a ground-truth for understanding the implications of different imaging protocols on the assessment of LV function using CT.

Using a model for normal LV function and a model for regional dysfunction of moderate size and severity, a set of parameters for the CPD registration algorithm was identified for our task. This was the first attempt at identifying the optimal set of CPD parameters for the quantification of human regional LV function using SQUEEZ. With these parameters, the accuracy of SQUEEZ for detecting programmed dysfunctional regions was measured. The accuracy of the strain estimate at the center of an abnormal region decreased as a function of decreasing size of the abnormal region; dysfunctional regions of 1 (28 mm) and 2/3 (19 mm) AHA segments as core diameter were captured with high accuracy, while the severity of dysfunction in the 1/2 (14 mm) and 1/3 (9.5 mm) AHA segments was underestimated. It should be noted that dysfunctional regions of 1/3 AHA segment and smaller are rarely detected by any current method in the clinical setting; however, as part of future work, more advanced approaches need to be developed to refine the SQUEEZ pipeline (modify CPD or use a different registration technique, modify mesh design) to facilitate the accurate detection of smaller localized wall motion abnormalities. Additionally, the effect of the spatial resolution of CT image acquisition on the accuracy of mesh registration and SQUEEZ needs to be investigated. Low dose CT scans can be spatially smoothed to increase SNR; the performance of SQUEEZ on smoothed data will determine the minimum dose achievable for obtaining local function with 4DCT.

### 4.1 Limitations

While the analytical phantom served as a ground-truth in the optimization and quantification of the spatial resolution limits of SQUEEZ for the detection of regional wall motion abnormalities, it does not exactly represent the texture change of the human LV during systolic contraction. Although a low-pass filter with a frequency response similar to that of a clinical CT scanner was applied to simulate the change in endocardial features, human LV systolic contraction in hearts with high ejection fraction lose endocardial features in a complex, non-random manner (for example, convergence of the papillary muscles and heart wall at end systole).

Currently, regional dysfunction is modeled in our phantom in a circular pattern, with the most severe dysfunction at the center and a smooth tapering off towards normal function radially outward. The model is limited by its symmetric shape which is likely adequate to model small regions of dysfunction; however, regional dysfunction occurs in various shapes, often in a non-isotropic manner, especially in severely abnormal hearts. While these abnormalities are simpler to detect, the accuracy of regional dysfunction quantification with non-isotropic spatial scales in *x, y,* and *z* may need to be investigated.

While the model offers a great degree of flexibility in programming regional dysfunction, it is currently limited by the lack of an accurate model of CT noise in the independent systolic phases for images with extremely low signal-to-noise ratio (i.e. images with doses < 0.2 mSv). Future work should address the effects of increasing CT noise and segmentation outliers on the computed RS_CT_ values so that the optimal smoothing can be applied and the lowest radiation dose limit estimated.

## 5. Conclusion

SQUEEZ is a user independent method for measuring endocardial regional shortening from 4DCT images acquired with routine clinical protocols. While SQUEEZ has previously been validated in pigs and canines, we performed the first analytical evaluation of SQUEEZ using a ground-truth human anthropomorphic LV phantom prescribed with physiological end systolic endocardial strains. The optimal registration parameters for our task were identified and the smallest lesion size that can be accurately captured using SQUEEZ was quantified. For an error tolerance of ± 5% from the programmed ground-truth, SQUEEZ with Coherent Point Drift motion tracking, can accurately measure strain in a dysfunctional region of 19 mm or greater in diameter. The threshold for *detection* of the existence of an abnormal region is between 14 mm and 19 mm.

## Appendix A Definition of dysfunctional patch as a 2D sigmoid surface

We defined our regions of dysfunction to be sigmoid in shape to simulate a smooth transition from the core of the dysfunctional region to the normal functioning endocardial tissue. The sigmoid function was defined as:

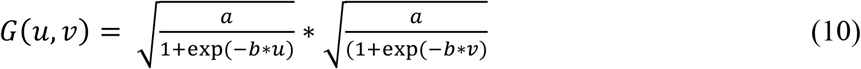

where *a* is the user defined % reduction in normal strain, *b* is the slope of the sigmoid function, and *u,v* are the planar coordinates defining a 71 x 71 grid from −3 to 4 in each direction; each user defined dysfunctional patch was normalized to this *u*,*v* grid. For the work reported in this paper, we fixed *b = 1.25* and *a* took on either 70% or 40% for simulating hypokinesia and subtle hypokinesia respectively.

To apply this smooth transition in strain, we parameterized the mesh points belonging to the dysfunctional patch in 3D space from *p(x,y,z)* to surface coordinates *s(l,z)*. By doing so, each point in the dysfunctional patch was given an index location based on its distance *l* along the endocardial surface from the user-input center of the dysfunctional patch and also on its slice location *z* relative to that of the patch center. Figure 7 shows three views of a user defined dysfunctional patch of the endocardium of radius 25 mm.

**Fig. 7.**
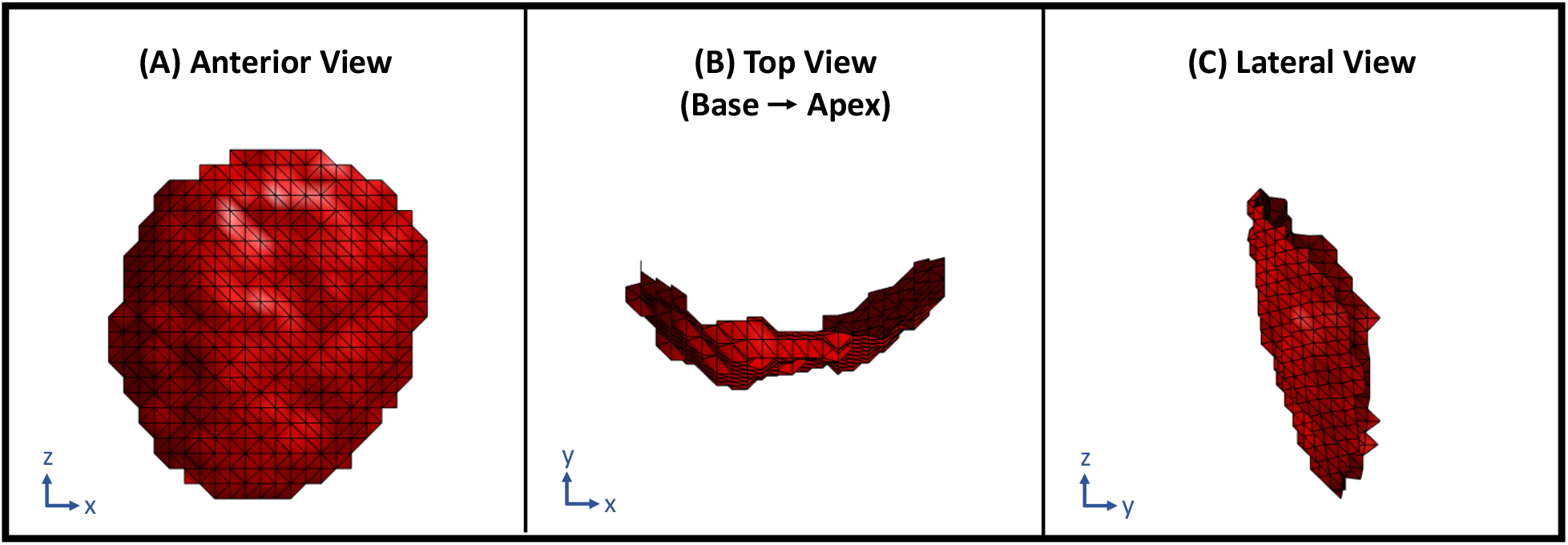
User defined LV endocardial patch of radius 25 mm to simulate regional dysfunction. (**A**) Anterior view of the patch with the septum to the left and the lateral wall to the right. (**B**) Top view of the patch looking from the base down into the apex with the inferior wall at 12 0’clock, lateral wall at 3, anterior wall at 6, and the septum at 9. (**C**) Lateral view of the patch with the anterior wall to the left and the inferior wall to the right.

The parameterization was performed in a slice-by-slice fashion; the number of rows in the 2D sigmoid matched the number of *z* slices in the dysfunctional patch. In this manner, each point on the dysfunctional patch was assigned a corresponding strain reduction factor defined by the sigmoid function. Points on slices closer to that of the user-defined center received a higher strain reduction than points farther away in *z*. Similarly, points in a particular slice closer to the user defined patch center along the surface in the circumferential direction received a higher strain reduction than points lying farther away. Figure 8A shows a single slice of points on the dysfunctional patch with the user-input patch center shown in red and Fig. 8B shows the 2D sigmoid function with a smooth transition from 70% reduction in strain at the center to ∼0% at the peripheral regions. The strain reduction factor assigned to each point in the dysfunctional patch was then multiplied with the ε_zz_ and the ε_cc_ components in Eq. 1-3 to create the desired region of dysfunction.

**Fig. 8.**
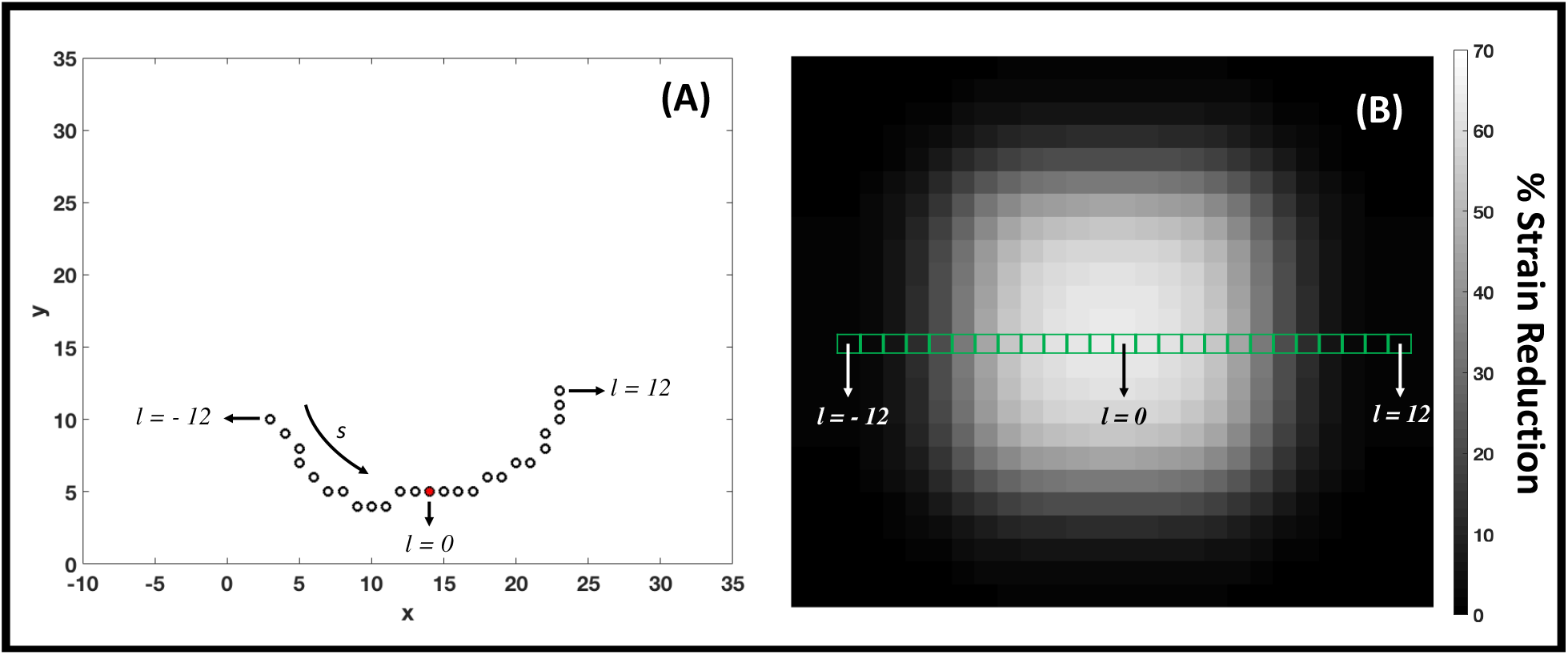
Application of 2D sigmoid function to ensure smooth transition in strain between the core of the dysfunctional region and the normal functioning endocardium. (**A**) Top view of a single *z* slice of points looking from the base down towards the apex with the inferior wall at 12 o’clock, lateral wall at 3, anterior wall at 6, and the septum at 9. The black circles represent points on the endocardial surface belonging to the dysfunctional patch with the user-input patch center highlighted in red. (**B**) 2D sigmoid function showing a smooth transition in strain from a peak 70% reduction at the core to no strain reduction at the peripheral regions. Each *z* slice of the dysfunctional patch corresponds to a row of the sigmoid function. When measured along the endocardial surface, points within a slice closer to the user-defined patch center received a higher reduction in strain than points lying farther away.

## Appendix B Frequency response estimation of the applied low-pass mesh filter: Comparison with the Standard reconstruction kernel of a clinical GE Revolution CT scanner

The low-pass filter, described in Sec. 2.3.3, was applied on the analytically displaced end systolic mesh points to simulate the natural endocardial texture change that occurs during systole. The texture change is a result of features collapsing during systole and the inability of the scanner to resolve these fine features due to its finite resolution (point-spread function). The filter required two user input parameters, the smoothing weight *α* and the number of iterations *n*. These values were chosen such that the modulation transfer function (MTF) of the filter was similar to that of the Standard reconstruction kernel of a 256 detector-row clinical scanner (Revolution CT, General Electric Medical Systems, Wisconsin). The Revolution CT was chosen for comparison purposes for the following two reasons: 1) the MTF of the Standard reconstruction kernel of this scanner has been measured previously^43^ and 2) it was the scanner available to us at the time to perform our own independent measurement of the MTF.

The experimental measurement of the MTF of the scanner was performed in a manner similar to that outlined by Cruz-Bastida et al., 2016^43^. A titanium bead of 0.25 mm in diameter was placed at the isocenter of the scanner and reconstructed (reconstruction diameter: 50 mm; kernel: Standard) at an in-plane resolution of 0.097 mm in *x* and *y*, with a slice thickness of 0.625 mm in *z*. 8 images with independent noise were acquired (tube current: 270 mA; kVp: 120 kV; gantry revolution time: 280 ms) and the MTF was measured by computing the discrete Fourier transform of the point spread function, obtained by averaging the 8 independent images of the bead. The estimated MTF was in good agreement with that measured by Cruz-Bastida et al., as seen in Fig. 9.

**Fig. 9.**
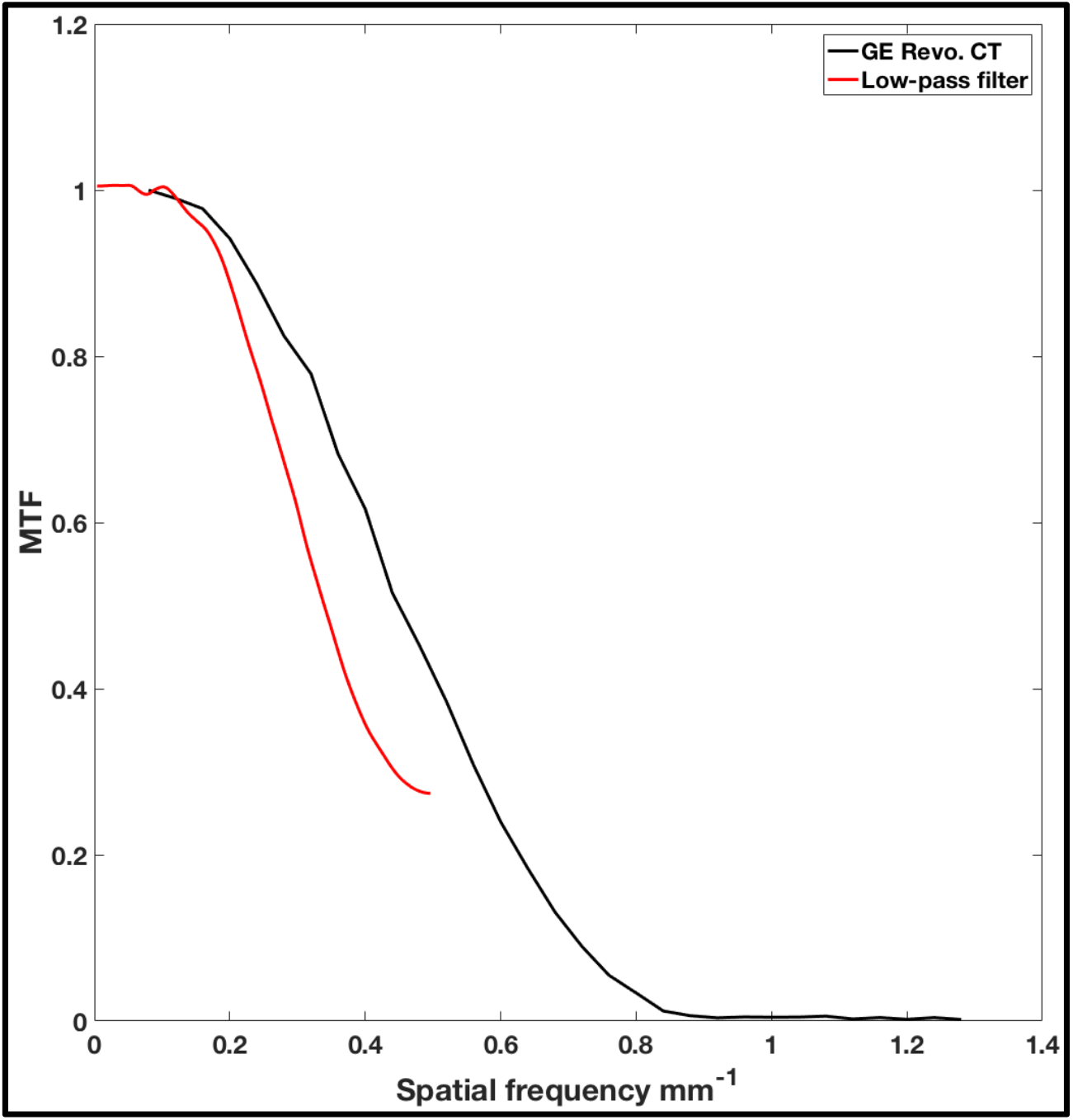
Modulation transfer function (MTF) of the GE Revolution CT Standard reconstruction kernel (black) and the low-pass mesh filter (red) applied on the analytically derived end systolic meshes.

The MTF of the low-pass filter, with the chosen values of *α = 0.4* and *n = 4*, was measured by creating 1000 images of 256 x 256 pixels of uniformly distributed white noise. A mesh was extracted with the defined 256 x 256 grid as the *x* and *y* coordinates and the connectivity matrix was obtained using the Delaunay triangulation algorithm. The intensity of the white noise at each pixel location was added as the *z* coordinate to the corresponding mesh point. The low-pass filter was independently applied to each of the 1000 meshes generated and the output of the filter in each case was converted back to an image by transferring the smoothed *z* values to pixel intensities at their corresponding locations on the image grid. The MTF was estimated by averaging the discrete Fourier transform of the 1000 filtered images. Figure 9 shows the MTF of the filter overlaid with the MTF of the scanner.

## Appendix C Estimating confidence intervals for endocardial edge position

Noise was added to the (*x,y,z*) coordinates of the analytically derived end systolic meshes prior to application of the registration algorithm for the following two reasons:

1. To break the ‘analytical coherence’ of mesh features between end diastole and end systole
2. To simulate the natural variability in endocardial position due to CT noise

The level of added noise was chosen within the range of the root-mean-square variance in 3D endocardial position as determined from Sec. C.3.

### C.1 Algorithm for Estimating Variability in Edge Location

If a stationary object was scanned in a CT scanner and independent images of the object were reconstructed with no view sharing, each reconstructed image would have independent noise, causing a degree of variability in the edge estimates. The method proposed to calculate the variability in edge location is outlined below:

1. Obtain multiple images *N* (>10), each with independent noise
2. Segment the object in each image
3. Sum all segmented images of the object
4. Calculate the number of pixels (voxels in 3D), *T*, in the transition range of the summed image (transition range defined by [1, N-1])
5. Detect the mean object from the summed image (defined by the object with pixel values >= N/2)
6. Calculate the perimeter (area in 3D) *P* of the edge of the mean object.
7. Edge variability *V = T/P*

The edge variability *V* is an estimate of the size of the uncertainty region within which the edge may be detected. It has the unit of length.

### C.2 Validation of Algorithm Proposed in Sec. C.1

The proposed method outlined in Sec. C.1 was validated by simulating a square with a programmed region of uncertainty within which the edge maybe detected under the influence of noise. The programmed ground-truth uncertainty was compared with the estimate obtained from the algorithm outlined in Sec. C.1.

From a clinical scan of a human LV (tube current: 530; kVp: 80 kV; gantry revolution time: 280 ms; reconstruction diameter: 200 mm; kernel: Standard; contrast agent: Visipaque 320 mg/ml, GE Healthcare, Chicago IL; contrast delivery: intravenous), an estimate of the gradient of the endocardial edge was determined for use in our simulation. The mean (μ) and standard deviation (σ) of HU within the contrast enhanced LV blood pool was measured to be 600 and 56 respectively and the user selected threshold for LV blood segmentation was set at 300 HU; thus 300 – 2*σ = 188 defines the HU limit for a 95% certainty in classification of the contrast enhanced blood pool from the myocardium. By setting the appropriate window level (300 HU) and window width (4*σ = 224 HU), it was observed that the HU transitioned from the blood pool to the myocardium over ∼2 mm for a 95% certainty in classification of each region. The edge gradient was spatially dependent; the gradient was more gradual near the papillary muscles and sharper at the base.

In an image of 512 x 512 pixels, a centrally located square object of side length 255 pixels with a uniform pixel intensity of 580 was created. The background of the image had a uniform intensity of 100. The intensity values were chosen to simulate that of the blood contrast agent in the LV (580 HU) and the myocardium (100 HU). For our simulation, the edge gradient from the square object to the background was linearly defined over 5 pixels (2.5 mm at 0.5 mm in-plane resolution); this enabled the edge gradient to be symmetric about a mean edge at a HU value of 340. The object (Fig. 10A) and a zoomed in view of the edge (Fig. 10B) are shown in Fig. 10.

**Fig. 10.**
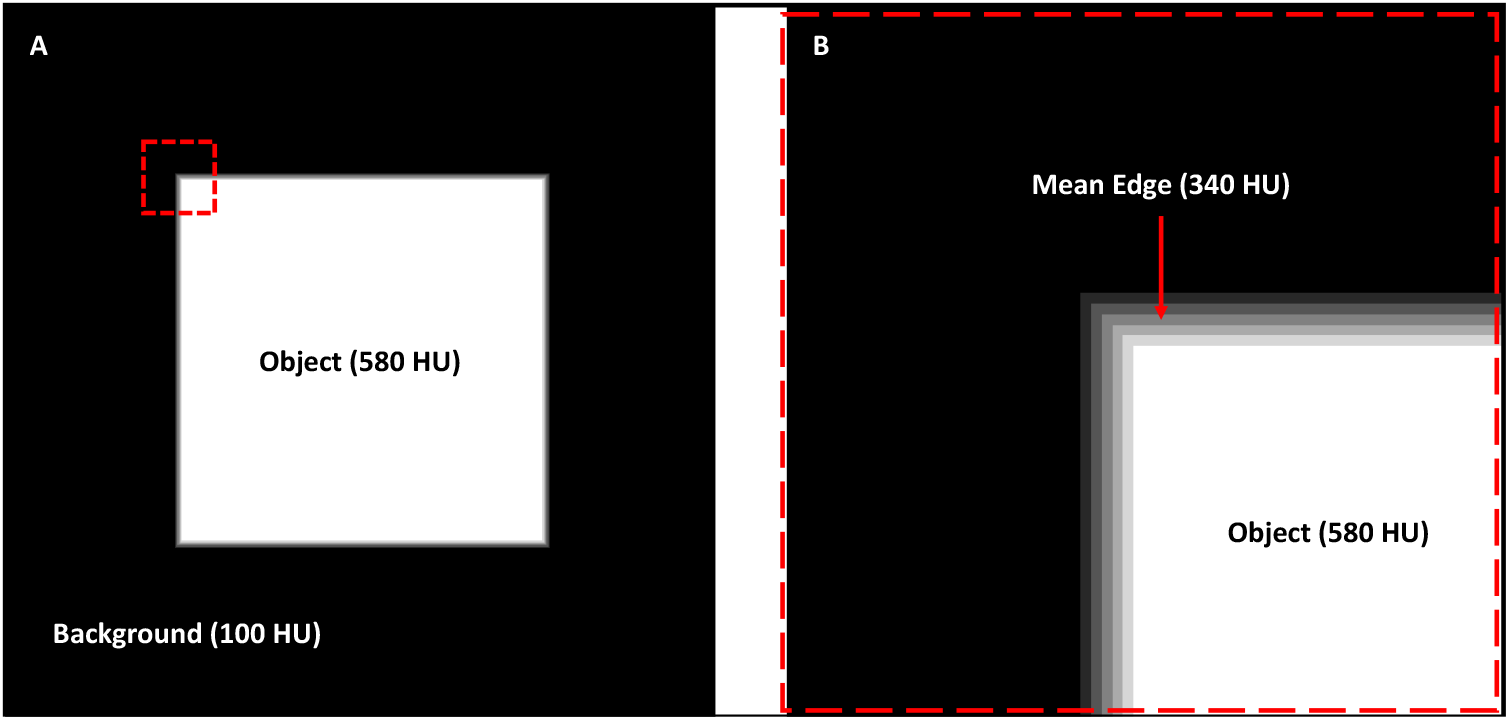
Simulation of precision estimation in endocardial edge detection. (**A**) Image of 512 x 512 pixels with a background intensity of 100 units (to simulate the myocardium HU) and a centrally located square object of size 255 x 255 pixels having a uniform intensity of 580 units (to simulate the blood contrast agent HU in the LV). The edge of the square object is defined by a linear gradient in intensity from the background to the object, occurring over a length of 5 pixels. (**B**) Zoomed in view of the object’s edge, corresponding to the dashed red box in (**A**), highlighting the gradient and the mean edge.

For a 95% certainty in classification of both the background and the square object, a uniform distribution of random noise in the range of ± 2σ (± 2*56), as determined from the clinical scan mentioned above, was added across the entire 512 x 512 image. The process was repeated 100 (*N*) times, yielding 100 images with independent noise and for each case, the square object was segmented by applying a threshold of 340 HU. For the above defined noise level and edge gradient, the programmed region of uncertainty in edge location was 3 pixels, and the measured estimate using the algorithm outlined in Sec. C.1 was 3.02 pixels.

As a second iteration of validation, with all other parameters kept constant, the edge gradient defining the transition in HU between the object and the background was changed to 11 pixels (5.5 mm at 0.5 mm in-plane resolution), which was representative of the edge gradient near the papillary muscles in our sample clinical LV scan. Again, 100 instances of independent noise were added to the image and the square object was segmented in each case by applying a threshold of 340 HU. The programmed region of uncertainty in edge location was 5 pixels, and the measured estimate using our algorithm was 4.99 pixels. In both simulations performed above, the estimates of variability in edge location obtained from the algorithm outlined in Sec. C.1 were in good agreement with the programmed ground-truth region of uncertainty in edge location.

### C.3 Precision Estimation of Edge Detection under Different Imaging Protocols

The end diastolic phase of a human LV endocardium derived from a clinical cardiac CT scan was 3D printed using a Formlabs 2 stereolithography printer (Formlabs Inc., Massachusetts) and the LV cavity was filled with an iodinated non-ionic radiocontrast agent (10% by volume solution of Visipaque 320, GE Healthcare, Chicago IL). The 3D printing material, which is a proprietary photopolymer resin (Formlabs Clear FLGPCL04), and the 10% iodinated contrast solution had HU values of 100 ± 9 and 502 ± 4 respectively. The printed phantom was imaged under 50 mA (9 mAs), 200 mA (36 mAs), and 400 mA (73 mAs) x-ray tube current settings (100 kVp; small focal spot) and for each tube current setting, 14 3D volumes were acquired and reconstructed (reconstruction diameter: 200 mm; kernel: Standard) with no view sharing such that each image had independent noise. The signal to noise ratios within a region of the LV blood pool for the three x-ray tube currents were measured to be 7.7 ± 0.2, 10.3 ± 0.9, and 12 ± 1.3. Precision estimates of edge location were obtained for each tube current setting using the algorithm outlined in Sec. C.1.

Figure 11 shows an axial slice of the phantom for each of the three tube current settings. The phantom was positioned in the scanner upright; hence, the axial views shown in Fig. 11 do not correspond to traditional axial views of a clinical scan with the heart in its native orientation. The variability *V* in the location of the edge was estimated using the algorithm outlined in Sec. C.1 to be 1.65, 0.9, and 0.65 mm (3.3, 1.8, and 1.3 pixels) for the 50, 200, and 400 mA tube currents respectively. Additionally, Fig. 12 shows three regions of an axial slice of the summation image (image obtained by adding the 14 independent LV segmentations) for each tube current setting (regions correspond to the dashed red, blue, and yellow boxes in Fig. 11). Line segments were drawn perpendicular to the endocardial surface that began at the high intensity region (pixel value = 14) and ended at the low intensity region (pixel value = 0), determined by visual inspection. The lengths of the line segments were used to quantify edge variability. These estimates were in agreement with the ones obtained using the algorithm outlined in Sec. C.1.

**Fig. 11.**
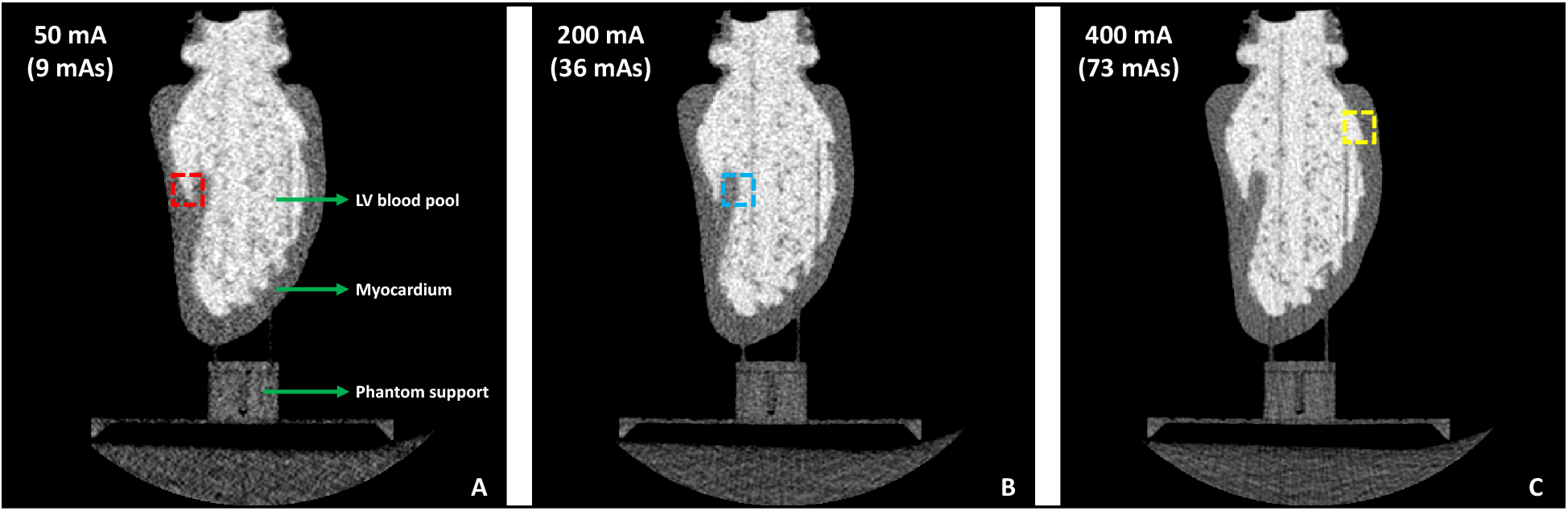
Axial slices of a 3D printed end diastolic human LV phantom imaged under different tube current settings. (**A**) 50 mA. (**B**) 200 mA. (**C**) 400 mA. For each tube current setting, the peak voltage was set to 100 kVp with a small focal spot size. The phantom was positioned in the scanner upright, hence these axial slices do not correspond to the axial slices of a clinical scan with the heart in its native orientation The dashed red, blue, and yellow boxes highlight the regions shown in the dashed boxes of the corresponding colors in Fig. 12.

**Fig. 12.**
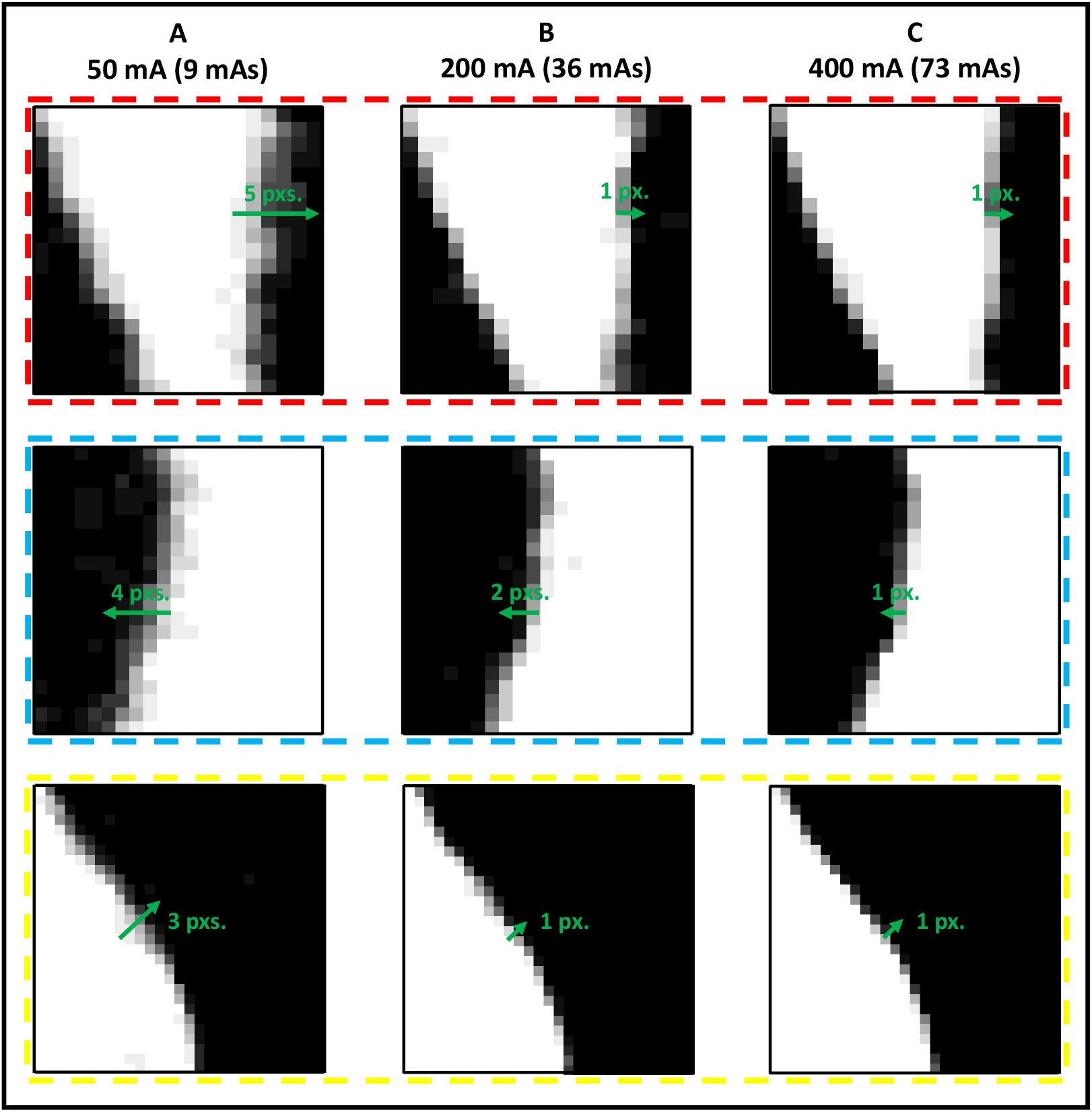
Variability in endocardial edge location at three highlighted zones of the segmented LV for different tube current settings. (**A**) 50 mA. (**B**) 200 mA. (**C**) 400 mA. The location of each zone is shown by the dashed boxes of the corresponding colors in Fig. 11. The images are of the summation volume obtained by adding the 14 independent segmentations of the LV blood pool for each tube current setting. White pixels are of intensity 14, black pixels are 0, and values in-between are represented by the various shades of gray. Line segments (in green) drawn perpendicular to the endocardial surface serve as visual aids for measuring the variability in edge locations.

The level of noise added to our analytically displaced end systolic mesh points (*x*,*y*,*z*) prior to the registration process was calculated from the precision estimates obtained above. The variability *V* for each tube current setting was divided by 2 to obtain an estimate of the variation about a ‘mean’. For example, for the 50 mA tube current setting, the edge had an uncertainty region of 1.65 mm; assuming the mean edge lies in the middle of this uncertainty region, the edge can be detected over a region of ± 0.83 mm. Similarly, for the 200 and 400 mA tube current settings, the regions of uncertainty were ± 0.45 mm and ± 0.33 mm respectively. Therefore, a mean value of ± 0.6 mm was added as noise to the end systolic mesh points to simulate the uncertainty in endocardial edge location.

## Disclosures

Dr. McVeigh holds founder shares in MR Interventions Inc. Dr. McVeigh receives research funding from GE Healthcare, Tendyne Holdings Inc., and Pacesetter Inc. No other author has any conflict.

## Acknowledgements

The authors would like to thank Dr. Marcus Y. Chen at the Laboratory of Cardiac Energetics, National Heart, Lung, and Blood Institute, National Institutes of Health (NIH), for providing us with the human CT data used in this work.

This work was funded in part by the NIH grants K01 HL143113 (FC), T32 HL105373 “Integrative Bioengineering of Heart, Vessels, and Blood” (GC), and R01 HL144678 (EM).

**Ashish Manohar** received a BE from the R.V. College of Engineering, India and an MS from UC San Diego, USA, both in mechanical engineering. He joined the Cardiovascular Imaging Laboratory (CViL) in 2017, where he is currently pursuing his doctoral degree in mechanical engineering. His research interests include cardiac function assessment from CT images, and understanding the spatial and temporal resolution limits of 4DCT in the estimation of regional wall motion abnormalities.

**Elliot McVeigh**, BSc in Physics and PhD in Medical Biophysics from University of Toronto. Faculty in BME and Radiology at Johns Hopkins from 1988-1999. PI at NIH/NHLBI from 2000-2007. From 2007 through 2015 he was Chairman of BME at JHU. Since 2015 he has been a Professor of Bioengineering, Radiology and Cardiology at UCSD. His laboratory focuses on creating novel MRI and CT techniques for diagnosis and guidance of therapy of heart disease.

